# Optogenetic cross-linking of the actin cytoskeleton using DARPins

**DOI:** 10.64898/2026.05.20.726659

**Authors:** Julia R. Ivanova, Ann-Cathrin Böttinger, Andreas Fink, Julia Köberle, Karim Ajmail, Amelie S. Benk, Dimitris Missirlis, Joachim P. Spatz

**Affiliations:** Max Planck Institute for Medical Research at Bildungscampus Heilbronn, Dept. of Cellular Biophysics, D-74076 Heilbronn, Germany; Max Planck School Matter to Life at Bildungscampus Heilbronn, D-74076 Heilbronn, Germany; Institute for Molecular Systems Engineering and Advanced Materials, Heidelberg University; postal address: INF 225, 69120 Heidelberg, Germany

**Author notes:** corresponding author: Dimitris Missirlis.

**Keywords:** DARPin, iLID, cytoskeleton, actin dynamics, lamellipodia, actin bundling, stress fibers

## Abstract

Precise tools to control actin filament organization and dynamics are essential for studying how the actin cytoskeleton regulates fundamental cell processes, such as morphological changes, migration and intracellular transport. Here, we present an optogenetic system for the reversible, spatiotemporal manipulation of actin cross-linking in living cells. Our approach employs actin-binding Designed Ankyrin Repeat Proteins (DARPins) fused to the improved Light-Induced Dimer (iLID) system, enabling actin cross-linking upon light-triggered dimerization. Following validation of cross-linking of bifunctional DARPins in reconstituted networks, the iLID system was integrated to create light-controlled associations inside cells. Optogenetic DARPin dimerization is rapid, reversible, and locally inducible. Activation of the DARPin-based actin cross-linkers revealed significant inhibition of cell traction forces when triggered throughout the cell. Our findings establish a versatile tool for investigating cytoskeletal dynamics with high spatial and temporal precision, paving the way for controlled manipulation of cellular architecture and mechanics.

**Graphical Abstract/Table of Contents Figure:** 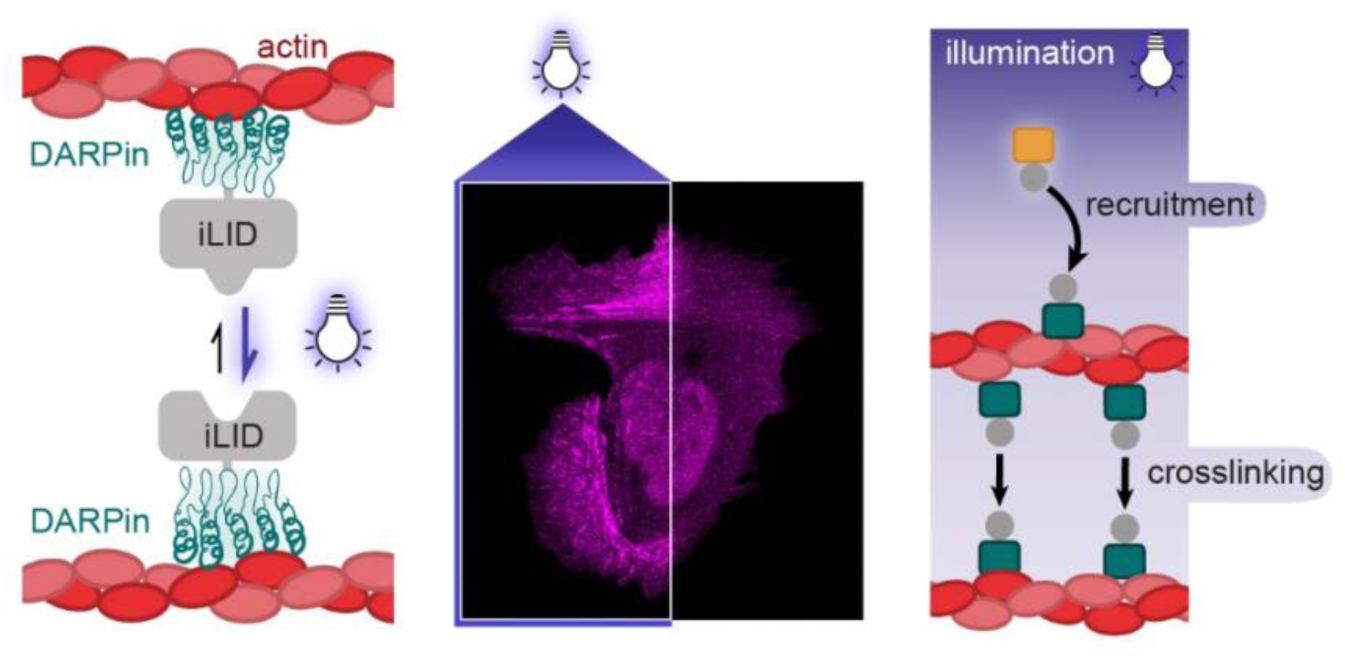

## Introduction

Actin filaments constitute a major building block of the cell cytoskeleton, the network of filamentous proteins that maintains cell shape and enables cell movements during migration or division. Filamentous actin (F-actin) is assembled from globular actin (G-actin) proteins into an asymmetrical structure that exhibits differential (de)polymerization kinetics on each of its ends. Inside the cell, F-actin assembly, disassembly and cross-linking are tightly regulated, thanks to a large number of actin binding proteins (ABPs), in order for the cytoskeleton to serve its dual function as a structural and a force-producing element.^1^ ABPs are involved in the nucleation, elongation, branching, bundling and severing of F-actin.^2^ In addition, motors of the myosin family are able to move along F-actin, or produce contractile forces when the motors are assembled in microfilaments.^3^ Our understanding of the structure and function of this complex interactome of proteins is still incomplete; new tools are necessary for studying and manipulating F-actin networks. Previous efforts have focused on fabrication of synthetic cross-linkers based on natural proteins, peptides and small molecules ^4–6^ for studying the behavior of reconstituted actin networks. However, F-actin cross-linkers for manipulation of the actin cytoskeleton in living cells, ideally with external spatiotemporal control and decoupling of physical from biochemical factors, are still lacking. While our manuscript was in preparation, an analogous, genetically-encoded cross-linker was reported,^7^ further highlighting the interest in manipulating F-actin organization in cells.

An attractive strategy to achieve precise, remotely-triggered manipulation of F-actin networks is to employ light and optogenetic systems. Such systems are in general composed of a light-responsive element, linked to a protein - or protein segment - that alters its structure upon illumination with specific wavelengths of light.^8, 9^ Light-triggered structural changes have been designed to result in affinity changes between molecules: an increase in affinity causes dimerization or oligomerization of the linked protein, whereas a decrease in affinity leads to their separation.^10–12^ For manipulation of F-actin, one ABP, which is linked to the optogenetic system, could be used to recruit a second protein to F-actin at the illuminated region. If the second protein is also an ABP, a reversible F-actin cross-linker would form through light control.

Here, we describe such a system based on DARPins (**D**esigned **A**nkyrin **R**epeat **P**rote**ins**) and the iLID (**i**mproved **L**ight-**I**nduced **D**imer) optogenetic system.^13^ DARPins are small (11-19 kDa) structurally-defined, synthetic proteins that can be selected for binding a target ligand through ribosome display.^14^ Various DARPins have been reported as targeting moieties for nanotechnology, as intracellular inhibitors, or as components of multivalent binders.^15–19^ We recently described the identification of several actin-binding DARPins with variable affinities, binding kinetics and intracellular distribution patterns, that can serve as effective F-actin labels.^20^ The iLID-nano optogenetic system was engineered from the light-oxygen-voltage 2 (LOV2) protein of *Avena Sativa*, through the incorporation of the bacterial SsrA peptide and further optimization: blue light induces a conformational switch in the protein that enhances affinity towards its ligand, SspB, by over 36-fold.^13^ By coupling actin-binding DARPins with the LOV2-SsrA or SspB proteins of the iLID nano system, we were able to control their dimerization inside living cells and affect actin-mediated traction force generation.

## Results & Discussion

### Bifunctional DARRins bundle actin filaments

We aimed to build an actin cross-linker that is controlled by light based on the iLID-nano system^13^ and the previously identified actin-binding DARPins.^20^ Prior to fabrication of the optogenetic tool, we validated the ability of divalent DARPins to cross-link purified actin filaments. To this end, a stable, homo-bifunctional DARPin was first synthesized. Single cysteines were introduced at the C-termini of DARPin 1784_A7, hereafter referred to as 1ABD (1 **A**ctin **B**inding **D**ARPin; **Table 1**). The introduced thiol groups on the cysteine were then covalently linked using a bis-maleimide ethylene glycol linker to produce the covalent DARPin dimer 1ABD-1ABD (**Figure 1A**). When incubated with fluorescently-labeled, phalloidin-stabilized actin filaments, 1ABD-1ABD homodimers bundled efficiently the actin filaments, similar to the actin-bundling protein fascin (**Figure 1B**). Accordingly, when co-encapsulated in giant unilamellar vesicles (GUVs) containing actin filaments,^21, 22^ the homodimer 1ABD-1ABD cross-linker induced bundling of actin filaments, in contrast to unorganized F-actin observed in absence of cross-linkers in GUVs (**Figure 1C**). Thus, bivalent actin-binding DARPins were able to cross-link purified actin filaments.

**Table 1.** DARPin constructs used in the present study. The DARPin unique identifiers and assigned nicknames, as well as the DARPIin-iLID constructs prepared are presented. The intracellular localization of the actin-binding DARPins when expressed with a fluorescent tag in living U2OS cells, as well as their recovery half-time on stress fibers as measured by FRAP, which reflects their actin-binding dynamics, are given.

| DARPin | DARPin nickname | DARPin-LOV2-SsrA domain | DARPin-mApple-SspB-nano domain | localization<sup>19</sup> | FRAP T<sub>1/2</sub><sup>19</sup> |
| --- | --- | --- | --- | --- | --- |
| 1784_A7 | 1ABD | 1ABD-SsrA | 1ABD-mApple-SspB | F-actin: lamellipodia & stress fibers | 0.33 ± 0.13 s |
| 2358_A11 | 2ABD | 2ABD-SsrA | 2ABD-mApple-SspB | F-actin: stress fibers | 65 ± 17 s |
| E3_5<sup>23</sup> | NBD | NBD-SsrA | NBD-mApple-SspB | cytoplasm | — |

**Figure 1.**
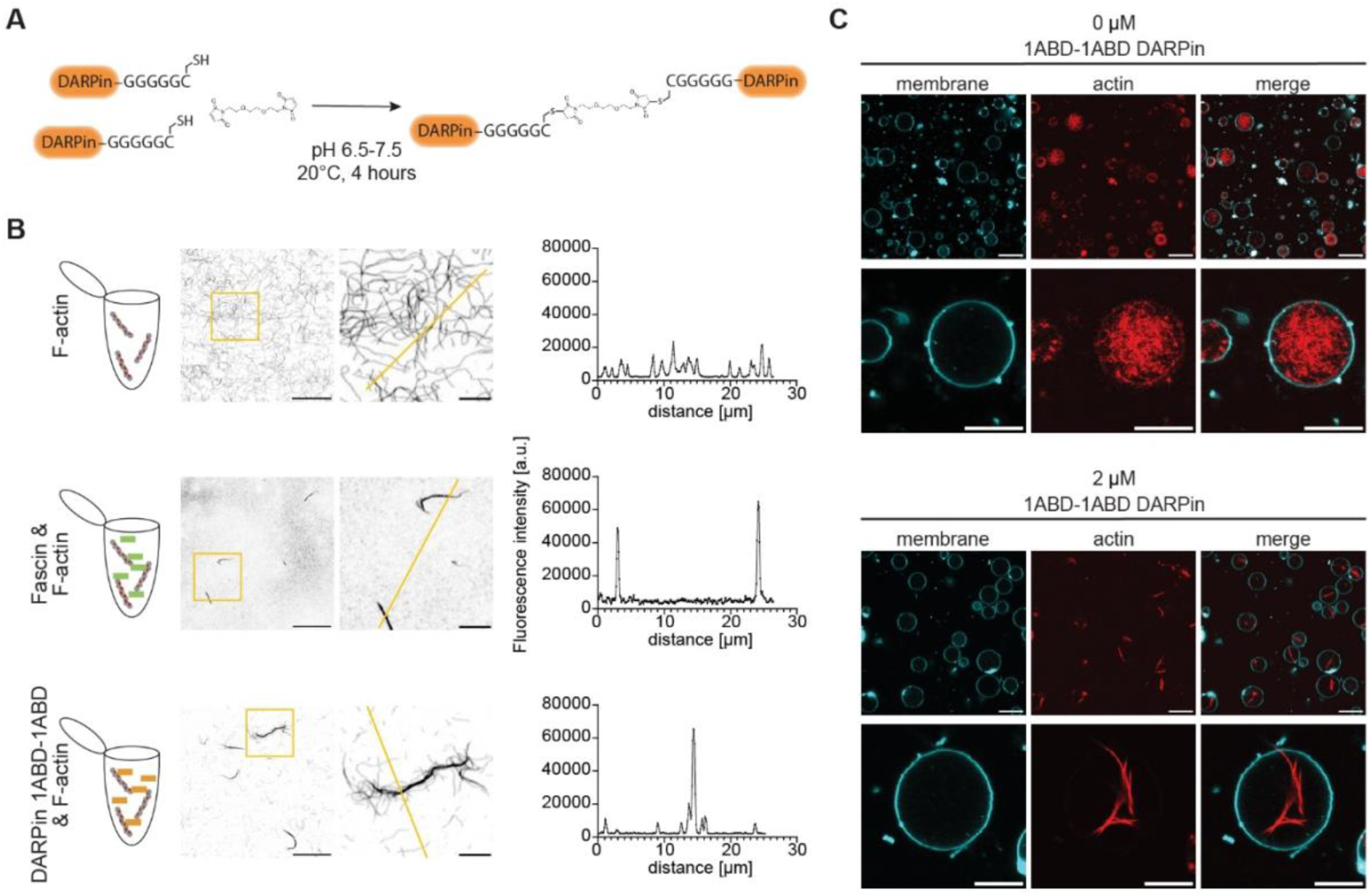
Covalent DARPin dimer 1ABD-1ABD bundles F-actin. (**A**) Purified DARPins 1ABD were C-terminally appended with a G_5_C peptide using a sortase-mediated reaction. They were then incubated with BM(PEG)3 that covalently linked two DARPins through the cysteine at their C-termini. (**B**) Confocal microscopy images (inverted) of TRITC-phalloidin-labeled F-actin (150 nM) that was incubated without a cross-linker, with fascin or with the covalent DARPin dimer 1ABD-1ABD. Scale bars: 20 µm overview, 5 µm zoomed region. Fluorescence intensity profiles were measured along the yellow line in zoomed images. (**C**) 5 µM F-actin was encapsulated in GUVs and labeled with SiR-actin (red). In presence of 2 µM of the covalent DARPin dimer 1ABD-1ABD bundling of F-actin was observed. The GUV membrane was labeled with Atto488-DOPE (cyan). Representative fluorescence confocal images are shown. Scale bar = 20 µm for overview images and 10 µm for images with single GUV.

### Optogenetic characterization of DARPin dimerization

To render DARPin dimerization reversible and controllable by light, DARPins containing the iLID components LOV2-SsrA and SspB-nano were designed, cloned and the plasmids were purified from *E. coli* lysates (**Table 1** & **Figure 2A**). DARPins were further labeled at the non-responsive SspB-nano unit with the fluorescent reporter protein mApple. Hereinafter, DARPin-mApple-SspB-nano constructs are referred to as DARPin-mApple-SspB and DARPin-LOV2-SsrA constructs are referred to as DARPin-SsrA (**Table 1**).

**Figure 2.**
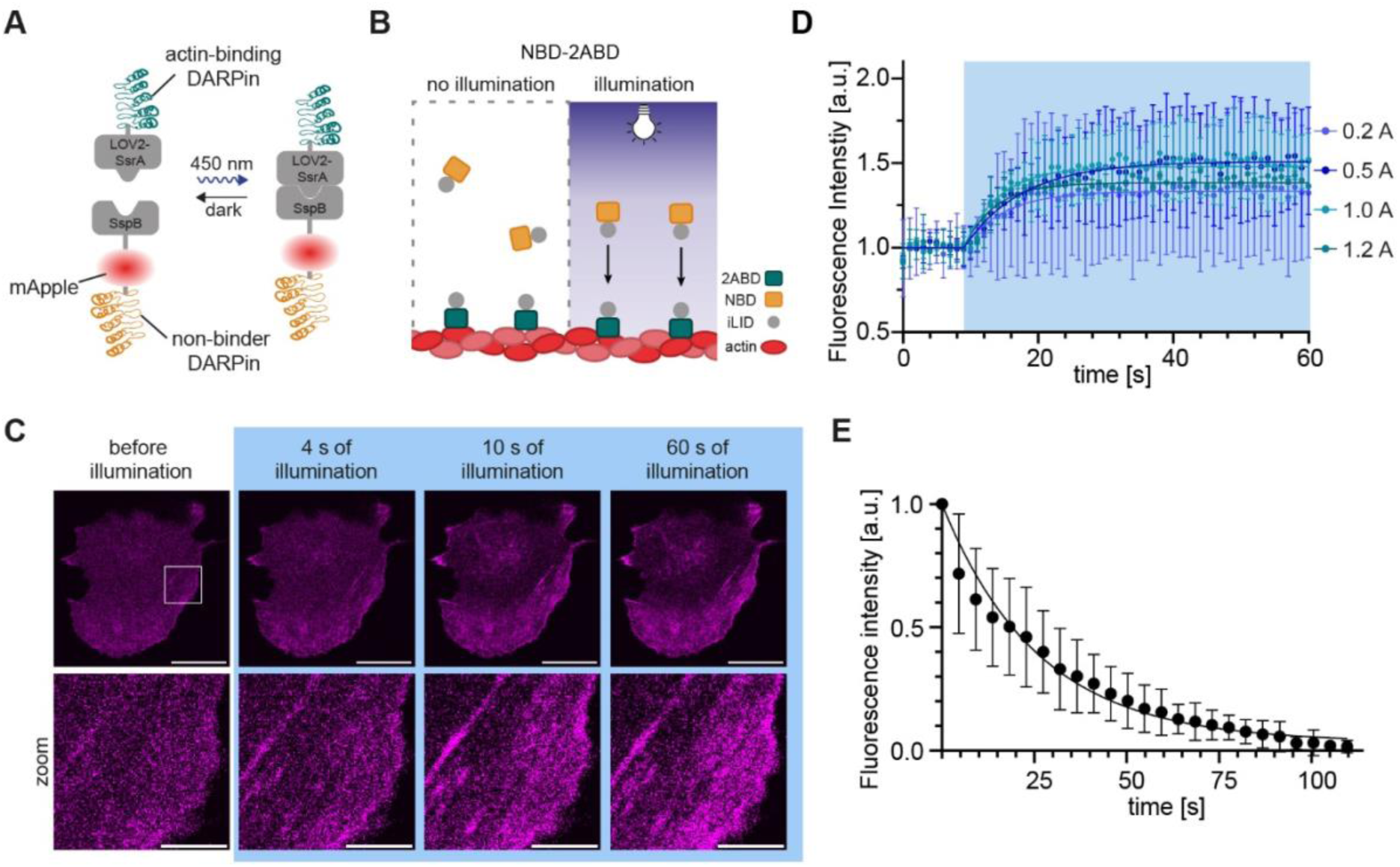
Dimerization dynamics of light-responsive iLID-DARPins. (**A**) The iLID system dimerizes during exposure with 455 nm light due to an increase in the affinity of LOV2-SsrA to SspB-nano from 4.7 µM in the absence of light to 130 nM during illumination with blue light. Accordingly, DARPins fused to the iLID domains dimerize under blue light. (**B**) A schematic representation of actin-bound 2ABD-SsrA proteins that dimerize with NBD-SspB proteins during illumination with blue light, causing a recruitment of fluorescent NBD-SspB to actin filaments. **(C)** Confocal microscopy images of live U2OS cells co-expressing 2ABD-SsrA and NBD-mApple-SspB (magenta). Cells were illuminated with 455 nm collimated LED light at 1.0 A. Enlarged images (zoom) corresponding to the region within the white rectangle are shown in the lower row. Scale bars for top row: 20 µm, bottom row: 5 µm. (**D**) Dimerization dynamics were monitored as the fluorescence increase over time in a stress fiber during illumination with 455 nm blue LED light at different power levels. (**E**) Dissociation dynamics of the iLID-DARPin dimer were measured by the fluorescence intensity decay from dissociating NBD-mApple-SspB on stress fibers upon stopping illumination. The fluorescence intensity was normalized between 1.0 and 0.0 and its decay was fitted with the following one-phase-decay equation: 𝑦 = (𝑦*_0_* − 𝑦_𝑚_) ∗ 𝑒(−𝑘𝑡) + 𝑦_𝑚_ where 𝑦*_0_* = *1*.*0*.

Among the actin-binding DARPins previously identified,^20^ we selected one with fast actin binding turnover (1ABD), one with slower dynamics (DARPin 2358_A11, hereafter referred to as 2ABD), as well as DARPin E3_5 with no known intracellular binding partner (non-binder DARPin: NBD) ^23^ (**Table 1**). The fluorescent NBD-SspB fusion served as a model protein for light-triggered recruitment to actin filaments and for kinetic studies (**Figure 2A,B**). DARPins 1ABD and 2ABD were previously shown to label F-actin in U2OS cells, albeit with distinct patterns: while 1ABD localized in more dynamic actin structures such as lamellipodia and filopodia, along with stress fibers, 2ABD accumulated only in stress fibers^20^.

When NBD-mApple-SspB and 2ABD-SsrA were co-expressed in U2OS cells, NBD-mApple-SspB exhibited homogeneous intracellular distribution in the dark, as expected since it was selected to have no binding partners (**Figure 2C**). However, upon blue light illumination with a 455 nm light-emitting diode (LED), NBD-mApple-SspB rapidly accumulated at F-actin structures (**Figure 2C**). The enrichment of the fluorescent DARPin construct on F-actin compared to the surrounding cytoplasm demonstrated the light-triggered association of NBD with the actin-bound 2ABD. The increase of fluorescence intensity at stress fibers over time was used to calculate the association half-time of the optogenetic DARPin system (**Figure 2D**). The association half-time ranged between 3.2 s and 6.0 s for the different LED driving currents used, which correspond to different light doses (**Figure 2D**). At a LED driving current of 0.2 A, the iLID-DARPin system associated with a half-time of 4.1 s (<14.3 s in 95% confidence interval (CI)); at 0.5 A the association half-time was 6.0 s (3.0 - 10.3 s in 95% CI), at 1.0 A it was 3.6 s (2.0 - 5.6 s in 95% CI) and at 1.2 A it was 3.2 s (2.4 - 4.1 s in 95% CI). Hence, the association kinetics were not significantly affected by the LED power in the examined range, suggesting that the light-induced conformational change of SsrA was not the rate-limiting step of the process. The conformational change of the blue-light-responsive LOV2 domain and the adjacent Jα helix, which allows for association between the SsrA and SspB domains, occurs in the millisecond time scale,^24, 25^ and robust association of the iLID system has been observed within seconds, even after a very short 10 ms pulse of light.^26^ Inside cells, the association kinetics depend on the diffusion of the protein of interest (SspB-fusion) to the illuminated complementary pair (SsrA-fusion), and therefore will also depend on the area and location of the illuminated ROI. Hence, it is not surprising that the kinetics we observed were faster than the previously reported recruitment of fluorescent proteins fused with the SspB-nano unit to locally activated LOV2-SsrA anchored in the plasma membrane (95.4 ± 18.1 s) or in mitochondria (21.2 ± 1.4 s),^27^ since in our experiments we illuminated the whole cell and the distance from any point in the cytoplasm to the nearest F-actin is shorter than to a small region of the plasma or mitochondrial membrane.

Next, we induced the localized, subcellular activation of the iLID nano system by using the laser of a confocal microscope as the light source for the same NBD-2ABD DARPin pair. We first validated that scanning the whole field of view using the 458 nm Argon laser line was sufficient to induce association of NBD-mApple-SspB with actin-bound 2ABD-SsrA (**Supp. Figure 1A**). The association between the two DARPins at the examined laser power intensities occurred rapidly, with half-lives between 1.5 and 1.9 seconds (**Supp Figure 1B,C**), similar to what was observed using the external LED illumination. When we activated half of the cell using the blue laser (458 nm) prior to imaging of NBD-mApple-SspB (561 nm), the accumulation of NBD-mApple-SspB on stress fibers occurred only on the illuminated side of the cell (**Supp. Figure 2**). Moreover, the association was reversible as evidenced by repeated association-dissociation cycles linked to light-dark cycles (**Supp. Figure 2**). When the blue laser illumination ceased, NBD-mApple-SspB returned to a homogeneous distribution inside cells, with a dissociation half-time of 18.1 s (16.1-20.3s in 95% CI) (**Figure 2E**). Overall, our results suggest that light can be used to recruit a protein fused with the SspB-nano peptide onto DARPin-decorated actin filaments in a rapid and reversible manner. This generalizable approach of triggering the artificial association of proteins with the actin cytoskeleton is reminiscent of a similar strategy actually used by cells to regulate protein activity in response to actin stress fiber assembly.^28^

### 3. Actin-binding DARPin dimerization inside cells

Contrary to 2ABD, which mainly accumulates on stress fibers, the more dynamic 1ABD localizes both at the cell edge in lamellipodia/filopodia and more centrally-located stress fibers. Hence, we selected DARPin 1ABD to induce homodimerization of actin-binding DARPins at the cell edge, inside a 9×9 µm² ROI. Accordingly, we co-expressed 1ABD-SsrA and 1ABD-mApple-SspB and used the laser scanning confocal microscope to repeatedly scan the subcellular ROI with blue laser light (458 nm). Intriguingly, 1ABD-mApple-SspB gradually accumulated at the illuminated spot (**Figure 3A,B**) reaching a steady-state, with an accumulation half-time of 79.6 s (70.2-90.2 s in 95% CI) (**Figure 3C**). The observed accumulation was not self-evident; the system was designed for light to induce dimerization of the actin-binding DARPins, and concomitant actin cross-linking at the ROI, and not recruit DARPins from other parts of the cell. We speculated that 1ABD-mApple-SspB moved from the surrounding cytoplasm to the ROI because 1) inside the ROI it had two binding partners, namely F-actin and 1ABD-SsrA, whereas outside the ROI, it could only bind F-actin, and 2) the light-induced dimer in the ROI exhibited a slower dissociation rate from the F-actin network, due to its bivalency (**Figure 3A**). Hence, the fluorescent DARPins would slowly accumulate in the ROI and exist in a dynamic equilibrium between F-actin-bound, DARPin-bound and free DARPin in monomer and homodimer form. The accumulation at the illuminated ROI was reversible; subsequent illumination of another ROI at a distance caused redistribution of the DARPin towards the newly irradiated area (**Figure 3B**).

**Figure 3.**
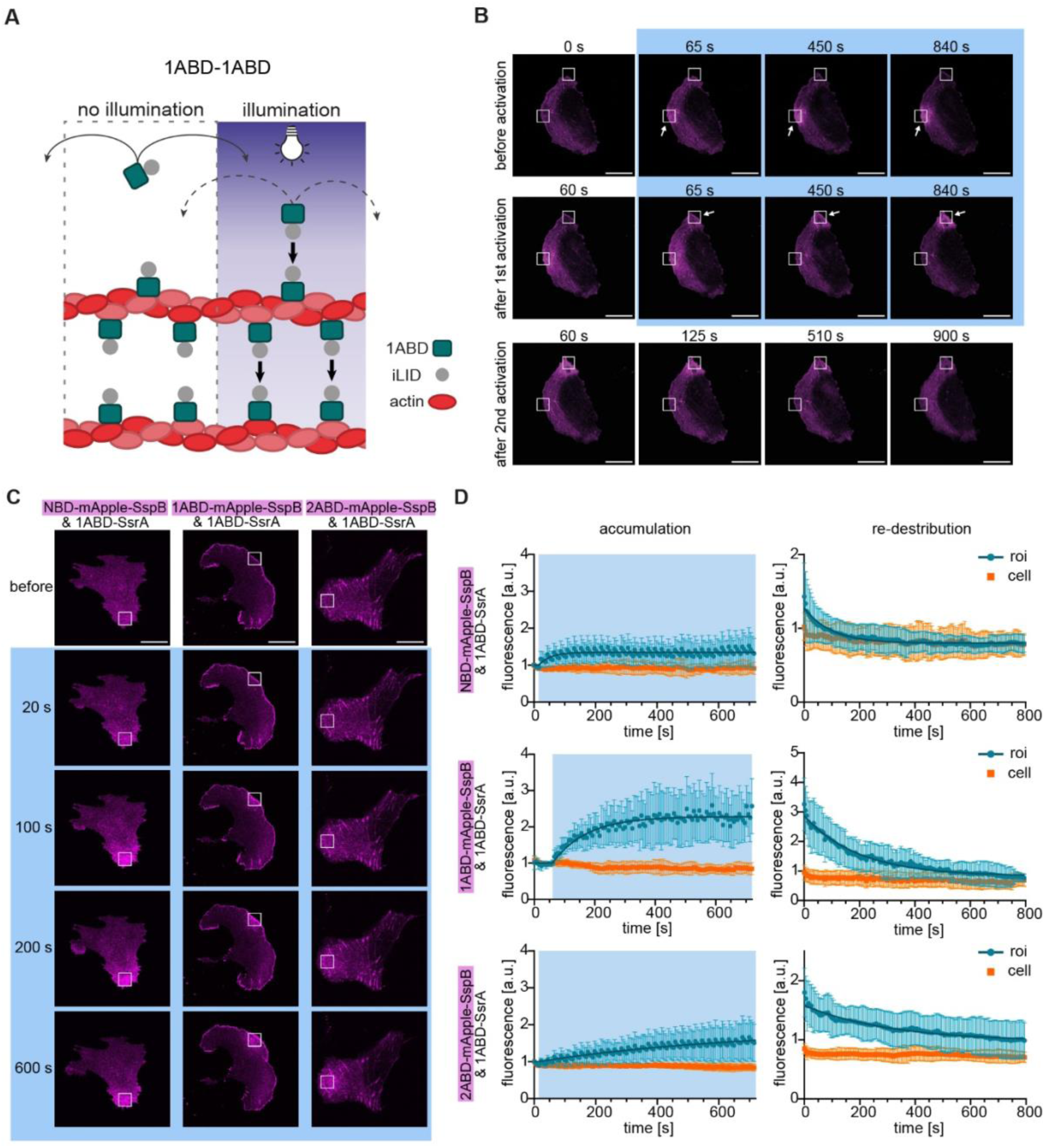
Dimerization dynamics of actin-binding DARRins in subcellular regions of living cells. (**A**) Schematic of actin-binding iLID-DARPins association under dark or illuminated conditions. In dark conditions, or in non-illuminated regions iLID-DARPin 1ABD can only bind to F-actin. In the illuminated regions, the iLID-DARPin 1ABD has two binding partners: besides actin, it can also bind another iLID-DARPin. (**B**) Live U2OS cell co-transfected with 1ABD-mApple-SspB (magenta) and 1ABD-SsrA. Fluorescence confocal images of a cell before, during (blue background) and after illumination with 455 nm light. Two different ROIs were subequently illuminated (marked with arrows) for 14 minutes, followed by monitoring of 1ABD-mApple-SspB fluorescence in absence of illumination. (**C**) Actin cross-linking was performed at a ROI, marked by the white outline, in cells co-expressing 1ABD-SsrA and the indicated, complementary DARPin-mApple-SspB unit. Confocal fluorescence microscopy images of typical cells with the different iLID-DARPin pairs before cross-linking and at different time points during cross-linking (blue background) are shown. (**D**) Normalized fluorescence intensity traces from the ROI (blue) and the whole cell (orange) as a function of time after 1) 455nm laser illumination, marked by the blue background (accumulation) and 2) termination of 455nm laser illumination (re-distribution). The traces were used to calculate the accumulation and re-distribution half-lives of the different DARPins. Scale bars = 20 µm.

The kinetics of accumulation at the ROI in this case did not represent the association kinetics between 1ABD-SsrA and 1ABD-mApple-SspB, which was expected to be similar to the one calculated for the NBD-2ABD pair (**Figure 2**). Instead, we hypothesized that the accumulation kinetics reflected the diffusion of unbound DARPins (monomers and dimers) in and out of the ROI, which are affected by the actin binding dynamics of DARPins. To test this, we exploited the distinct actin-binding dynamics of 1ABD and 2ABD. When unbound, both DARPins were expected to have similar diffusion in the cytoplasm due to comparable size and shape; however, 2ABD has a much slower turnover from F-actin compared to 1ABD as previously measured by FRAP (half-recovery time of τ_1/2_ = 65 ± 17 s for 2ABD compared to τ_1/2_ = 0.33 ± 0.13 s for 1ABD).^20^ Thus, intracellular movement of 2ABD was expected to be slower compared to 1ABD, since the former would be bound to the cytoskeleton for a longer time. Indeed, when the accumulation of NBD-mApple-SspB, 1ABD-mApple-SspB and 2ABD-mApple-SspB to a subcellular ROI with photo-activated 1ABD-SsrA were compared under the same conditions, the half-time of accumulation was significantly higher for 2ABD (454.1s; 321.6 - 733.1 s in 95% CI) compared to either 1ABD-mApple-SspB (78.4 s; 64.9 - 94.1 s in 95% CI) or NBD-mApple-SspB (26.8 s; 17.1 - 38.3 s in 95% CI), which has no binding partners in the cytoplasm (**Figure 3C,D**). This indicated that the rate-limiting step for accumulation at the activated region was the transport of DARPins inside the cells, which was slower for the DARPin with the slowest actin-binding turnover. Upon ceasing to illuminate the ROI, the DARPins redistributed in the cells, albeit with different kinetics: NBD-mApple-SspB redistributed faster (67.6 s; 56.8-80.6 s in 95% CI), than 1ABD-mApple-SspB (137.6 s; 117.0 - 164.2 s in 95% CI) and 2ABD-mApple-SspB distributed most slowly (216.8 s; 160.1 - 317.9 s in 95% CI).

In sum, our data indicates that iLID-mediated DARPin dimerization alters their subcellular binding dynamics, which are linked to changes in cytoskeleton binding, and leads to accumulation at activated regions.

### 4. Actin distribution while cross-linking inside cells

In order to examine potential changes of the local actin concentration at illuminated regions, F-actin was labeled independently from DARPins by co-expressing Lifeact-HaloTag and incubating cells with Halo-Ligand-AlexaFluor647 during activation. The fluorescence intensity of Lifeact within the illuminated ROIs did not change with time, indicating that the concentration of F-actin remained constant within the duration of the experiment (**Figure 4A,B**). Quantification of the ratio of AlexaFluor647 fluorescence signal inside the ROI to that of the whole cell did not change after 20 minutes of illumination (**Figure 4C,D**). In contrast, the fluorescence signal of DARPins increased within this duration (**Figure 4C,D**), as shown before (**Figure 3**). Similar results were obtained for both the 1ABD-1ABD and 2ABD-2ABD pairs (**Figure 4**). These results indicated that DARPin-mediated cross-linking of actin filaments does not affect the concentration of these filaments within the illuminated cell regions.

**Figure 4:**
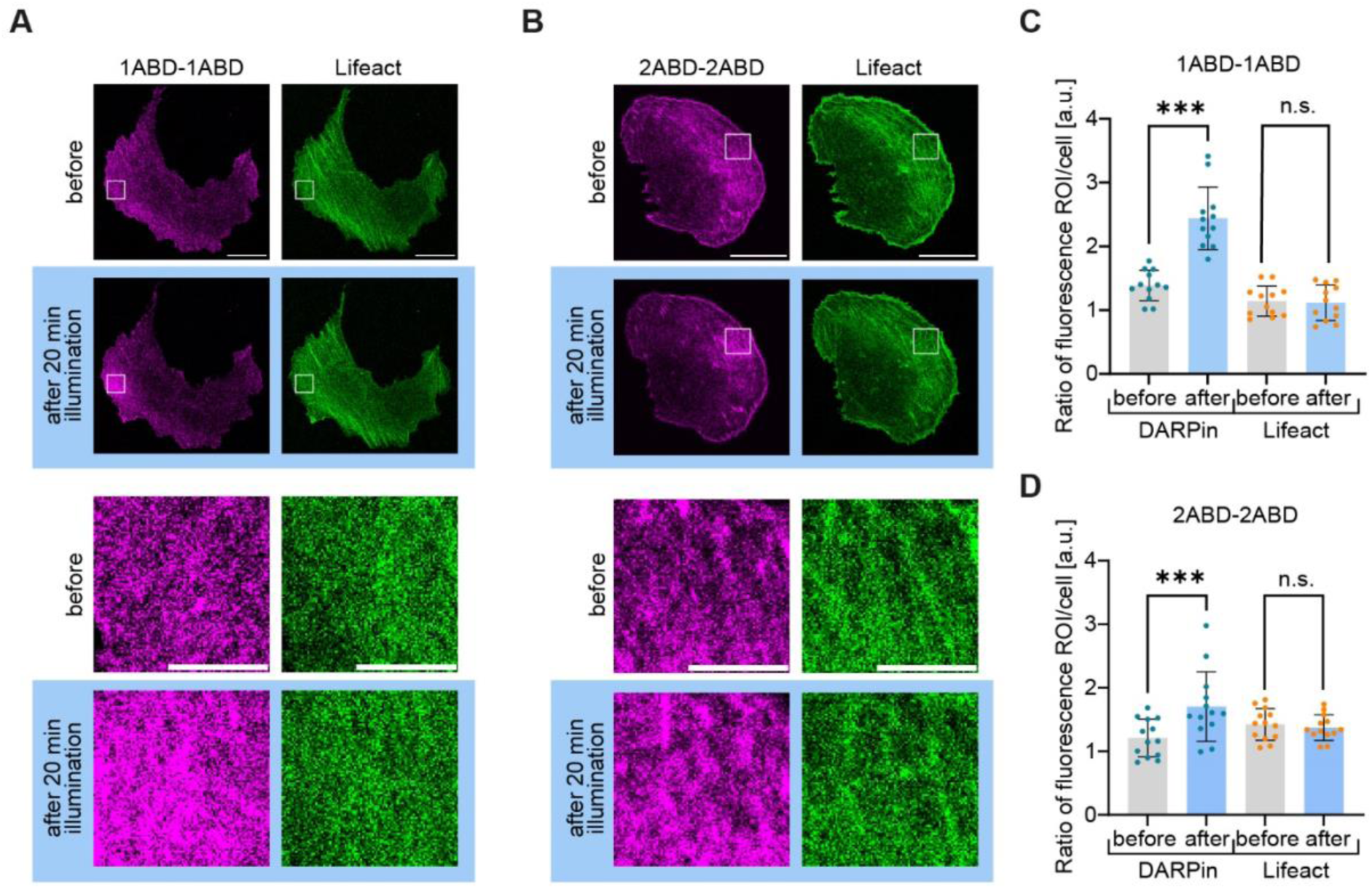
Light-controlled subcellular accumulation of actin cross-linker does not lead to actin accumulation. (**A,B**) Cells co-expressing Lifeact (green) with iLID-DARPin pairs 1ABD-1ABD (**A**) or 2ABD-2ABD (**B**) (magenta) were illuminated with 455 nm laser light inside a 9×9 µm ROI (white rectangle) during 20 minutes. Confocal microscopy mages of Lifeact and the corresponding DARPin-mApple-SspB before and after the 20-minute scanning are presented; enlarged images of the ROIs (white outlines) are shown below the whole cell images. Scale bars whole cell: 20 µm, zoomed region: 5 µm. (**C,D**) The fluorescence intensity ratio between the ROI and the whole cell increased after 20 minutes of illumination for the DARPin-mApple-SspB construct A3 (**C**) and A22 (**D**), but not for Lifeact-labeled actin.

### 5. Lamellipodia dynamics during enhanced actin cross-linking

Next, we explored the functional effect of light-controlled actin cross-linking in living cells. We first focused on the protrusion and retraction dynamics of lamellipodia, the actin-rich structures at the cell edge, that are involved in cell exploration of their surroundings and migration.^29^ The F-actin network in lamellipodia is highly dynamic with constant actin monomer addition at nucleated filaments at the cell edge and movement of the whole network towards the cell interior, known as actin retrograde flow; the ARP2/3 complex, which nucleates daughter filaments at an angle from existing ones, is essential for lamellipodia formation.^30^ The architecture of F-actin in lamellipodia can be altered in response to the concentrations of actin-binding and actin-regulating proteins,^31^ as well as force.^32^ Therefore, we hypothesized that optogenetic actin cross-linking might additionally affect lamellipodia dynamics.

In order to target lamellipodia, the iLID-DARPin pair 1ABD-1ABD was selected, since it efficiently accumulates at these dynamic structures.^20^ Regions at the cell edge that exhibited the characteristic high density of F-actin typical of lamellipodia were illuminated and the cell edge was monitored within the ROI during blue light illumination (**Figure 5A,B & Supp. Figure 3**). The change in cell area within the ROI, which reflected the protrusion versus retraction of the cell edge, did not exhibit a preferred trend: in some regions the area increased indicating cell protrusion, while in others the area decreased corresponding to cell retraction (**Figure 5C**). The net average of all samples was not significantly different from zero, when calculated over a period of 5, 10 or 15 minutes. The same experiment was additionally performed at a control (non-illuminated) region of the same cell, with the same outcome (**Figure 5C**).

**Figure 5.**
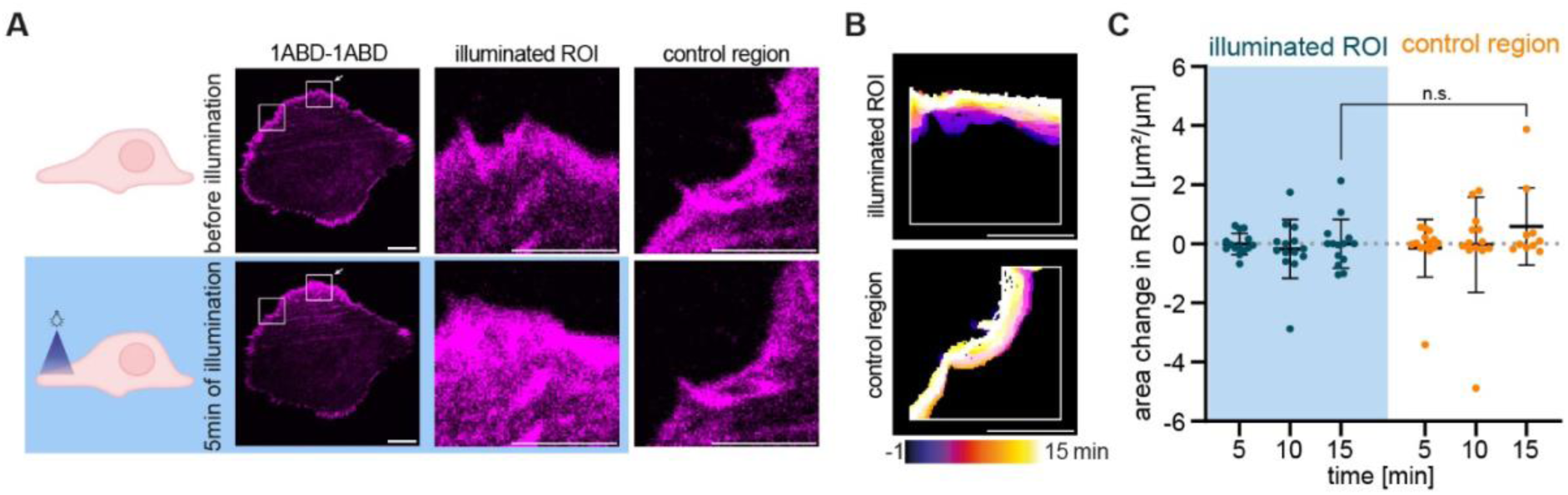
Functional effect of actin cross-linking on lamellipodia of living cells. (**A**) Actin was cross-linked in a ROI (white rectangle marked with arrow) enclosing part of the lamellipodium in cells expressing the iLID-DARPin pair 1ABD-1ABD. A non-cross-linked region of the same size, also including the lamellipodium (white rectangle without arrow) was used for comparison. Confocal microscopy images of the whole cell (scale bar: 10 μm) and enlarged regions of the illuminated and control ROIs (scale bar: 5 μm), before and 5 minutes after actin cross-linking are shown. (**B**) The cell edge outlines for both regions were monitored over time and plotted with a temporal color code. (**C**) The area within the ROI was calculated and normalized to the length of the lamellipodium in the ROI before and after 5, 10 and 15 minutes of illumination. The change in area for each time point was then calculated and plotted for the illuminated ROI (blue background) and the control region (Mean and standard deviation; n=10-14 cells; N=3 independent experiments).

Our results showed that optogenetically-mediated actin cross-linking at lamellipodia did not bias lamellipodia dynamics towards protrusion or retraction under the tested conditions. The lack of an effect on protrusion dynamics is likely due to insufficient cross-linking despite induced dimerization, since the dissociation of DARPin 1ABD is rapid. As the DARPin 2ABD did not localize to lamellipodia to test this, we turned our attention to the effect of cross-linking more stable actin-structures using the 2ABD-2ABD DARPin pair.

### 6. Cross-linking of actin filaments reduces traction forces

We examined the effect of F-actin cross-linking on the transmission of actomyosin contractile forces to their substrate and performed traction force microscopy under light-controlled cross-linking (**Figure 6A,B**). Previous studies on the role of endogenous actin cross-linkers, and in particular α-actinins, have reported contradictory results. In some reports, depletion of α-actinins enhanced traction forces,^33, 34^ while in others a reduction was observed.^35, 36^ Different hypotheses based on altered friction of actin fibers during myosin-induced sliding have been formulated to explain the data;^37^ however, this remains an open question, which is further confounded by potential changes in expression level of myosins,^38^ as well as the role and presence of α-actinins not only in stress fibers, but also at focal adhesions, which are the multi-protein complexes that mechanically link and transmit the intracellular forces from the cytoskeleton to the extracellular space.^39^

**Figure 6.**
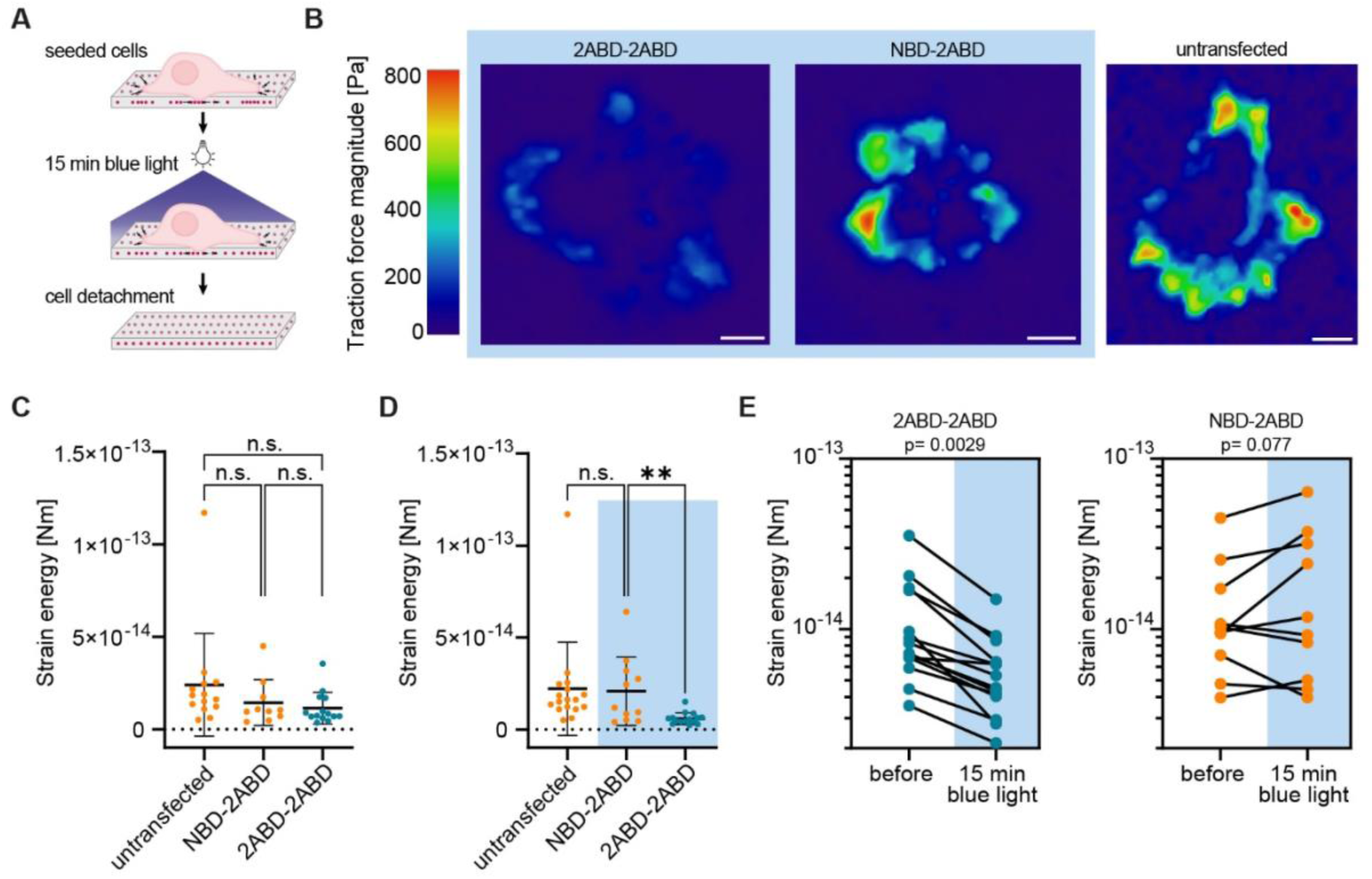
Actin cross-linking reduces traction forces of living cells. **(A)** Schematic of traction force microscopy experimental setup: the displacement of fluorescent beads inside elastic PAA hydrogels as a result of cellular traction was determined by comparing images before and after light illumination (15 minutes; 455 nm), as well as following cell detachment to determine relaxed positions. Cells expressing the iLID-DARPin pairs 2ABD-2ABD (actin cross-linker) and NBD-2ABD (control construct) were tested and compared to untransfected, control cells. **(B)** Representative traction force fields of cells expressing 2ABD-2ABD or NBD-2ABD immediately after 15 minutes of illumination as well as non-illuminated, untransfected cells (scale bar: 20 µm). **(C)** Calculated strain energy of cells expressing 2ABD-2ABD, NBD-2ABD and untransfected cells prior to illumination with blue light. **(D)** Strain energy of illuminated cells (15 minutes; 455 nm) expressing 2ABD-2ABD, NBD-2ABD; data for non-illuminated, untransfected cells are teh same as in (**C**) and added for comparison. **(E)** Comparison of strain energies for cells expressing 2ABD-2ABD or NBD-2ABD before and after 15 minutes of illumination with blue (455 nm) light. **(C-E)** Number of cells (n) and independent experiments: 2ABD-2ABD n=14-15 and N=3; NBD-2ABD n=10-11 and N=4; Untransfected cells n=13 and N=3.

In order to target DARPins at stress fibers and create optically-controlled cross-links between the actin filaments, the 2ABD-2ABD iLID-DARPin pair was used, along with NBD-2ABD as a control. Transfected cells on fibronectin-coated polyacrylamide (PAA) substrates with nominal Young’s modulus of 16.7 kPa^40^ exhibited similar strain energies compared to untransfected controls under dark conditions, demonstrating that the mere expression of the DARPin pairs did not impair actomyosin contractility (**Figure 6C**). After illumination of cells with 455 nm LED light for 15 minutes, cells expressing the 2ABD-2ABD pair showed a consistent decrease in traction forces as expressed here by the total strain energy (**Figure 6D,E**). The effect was specific to the actin cross-linking DARPin pair, as activated NBD-2ABD did not show any consistent trend in traction force magnitude (**Figure 6E**). Our data therefore suggest that actin cross-linking using a synthetic, reversible system without other known targets, leads to a decrease in cell traction forces.

Overall, we have introduced a novel, modular tool that we believe will find many applications and contribute towards addressing questions related to the F-actin structure-function relationship. While our manuscript was in preparation, an analogous, genetically-encoded cross-linker was reported,^7^ further highlighting the interest in manipulating F-actin organization in cells.

## Conclusions

Here we present an optogenetic actin binding or actin cross-linker approach based on the iLID-nano system and the actin-binding DARPins we previously identified. The variations in actin-binding dynamics among the different DARPins provided insight into the mechanism underlying their accumulation at activated regions; moreover, these variations provided a simple handle in modulating the degree and strength of induced protein links. A first attempt to modulate the protrusion rates of the highly dynamic actin-rich lamellipodia did not reveal a measurable effect most likely due to insufficient cross-linking; on the other hand, light-induced actin cross-linking over the whole cell resulted in a specific reduction of traction forces, highlighting the effectiveness of the proposed tool. In sum, our work demonstrates the applicability of this tool to precisely spatiotemporally control the dimerization of actin-binding DARPins, excluding complications that could arise by use of naturally-occurring cross-linkers and intermediary signaling.

## Experimental Section

### Materials

Lifeact-Halo plasmid and Halo-ligand coupled to Alexa Fluor647 were kindly gifted from the lab of Kai Johnsson (MPI for Medical Research, Heidelberg). All other materials were acquired from Merck, unless otherwise noted.

### Cloning

For mammalian expression of DARPins with a C-terminal mEGFP-tag, the plasmid mEGFP-N1 (a gift from Michael Davidson, Addgene plasmid # 54767; http://n2t.net/addgene:54767 ; RRID:Addgene_54767) was modified by NEB Builder high-fidelity (HiFi) DNA assembly. For this, the PCR primers 5’-GTGAGCAAGGGCGAGGAGCTGTTC-3’ and 5’-CTCGAGATCTGAGTCCGGTAGCGCTAG-5’ were used for plasmid linearization. A N-terminal Kozak sequence for mammalian expression, two restriction sites BamHI and HindIII for DARPin insertion to the multiple cloning site and a (G4S)2-linker between the HindIII restriction site and C-terminal mEGFP (monomeric enhanced green fluorescent protein) were inserted using the following dsDNA insert: 5’-CTACCGGACTCAGATCTCGGCCACCATGGGATCCGACCTGAAGCTTAATGGTGGCGGTGGCTCTGGCGGTGGTGGCAGCGTGAGCAAGGGCGAGGAGCTG-3’.

DARPins were inserted into the modified vector mEGFP-N1mod by restriction digest using the newly generated restriction sites.

In order to create DARPins that dimerize upon blue light illumination, domains of the iLID-nano system replaced mEGFP at the C-terminus of the DARPins by restriction digest. Plasmids containing genes for the LOV2-SsrA or SspB nano domain were a kind gift from Yan Jie (pmApple-TalinN10-GSGS-linker-LOVSsrA and SspB-GGSGGS linker-talinc11-mEmerald were provided by Samuel Barnett). The genes for the iLID domains were appended with restriction sites to replace mEGFP and a stop codon by PCR. For the PCR of the Lov2-SsrA construct the forward primer 5’-TGAAAGCTTGGGGAGTTTCTGGCAACCAC-3’ and the reverse primer 5’-AGGCGCGGCCGCTTAAAAGTAATTTTCGTC-3’ were used. For the amplification of the SspB gene the forward primer 5’-GCTAAGCTT-GAATTCAGCTCCCCGAAACGC-3’ and the reverse primer 5’-ATTGCGGCCGCTTAACCAATATTCAGCTCGTCATAGATTTCTTC-3’ were used.

For tracking, the fluorescent protein mApple was inserted together with a 6xHis-tag and flexible GSG linkers between the DARPin and SspB domain by Gibson Assembly. A PCR was performed to create a 5’ end at the beginning of the SspB gene and the 3’ end after the DARPin that is homologous to the ends of the mApple gene insert. For this, the primer 5’-GAATTCAGCTCCCCGAAAC-3’ was used as the forward primer. The reverse primer for the non-binder DARPins was 5’-TTGCAGGATTTCAGCCAG-3’ and for the actin-binding DARRin 5’-AGCAGCTTTCTGCAGAAC-3’.

Overhanging regions of the mApple gene insert were created with a forward primer containing at a region that is homologous to the DARPin, the GSG-6xHis-GSG sequence and a region homologous to mApple. The forward primer was 5’-ACCTGGCTGAAATCCTGCAAGGATCCGGACATCACCATCACCATCACGGCAGCGGTGTGAGCAAGGGCGAGGAG-3’ for the non-binder DARRin and 5’-AAGTTCTGCAGAAAGCTGCTGGATCCGGACATCACCATCACCATCACGGCAGCGGTGTGAGCAAGGGCGAGGAG-3’ for the actin-binding DARRins. The reverse primer was 5’-CGTTTCGGGGAGCTGAATTCGCCTGAACCCTTGTACAGCTCGTCCATG-3’ and contained a homologous region to SspB, the GSG linker and a homologous region to the 3’ end of mApple. The resulting gene construct was DARPin-GSG-6xHis-GSG-mApple-GSGSspB. Sanger Sequencing of the gene insert was performed by MicroSynth GmbH to validate the assembly of the desired product with the CMV-forward primer and a custom reverse primer: 5‘-GGCTGATTATGATCTAGAGTCGCG-3‘.

### Protein Purification

The sortase A pentamutant (eSrtA) in plasmid pET29 was a gift from David Liu (Addgene plasmid # 75144; http://n2t.net/addgene:75144 ; RRID:Addgene_75144).^41^ Purified DARPins with a C-terminal LPETGG-6xHis tag were processed using a sortase-based enzymatic reaction and a following Michael-addition to create covalent DARPin dimers. The sortase and DARPins were purified as previously described^20^.

### Covalent DARPin dimers

Purified DARPins were labelled C-terminally by sortase reaction with a GGGGGC peptide using a sortase-based enzymatic reaction. For this, DARPins expressed with a C-terminal LPETGG-6xHis (50 µM) tag were mixed with His-tagged sortase A pentamutant (2.5 µM) and the peptide GGGGGC (250 µM) (PSL peptide specialty laboratories GmbH, ID # 2358-12-20) in the sortase reaction buffer at 37°C for 30 minutes. Thereby, the DARPins C-terminal GG-6xHis peptide was replaced by GGGGGC. DARPins were separated from unreacted peptide and the buffer was replaced with phosphate buffer (48 mM K2HPO4, 4.5 mM KH2PO4, 14 mM NaH2PO4 in distilled water at pH 7.2) by ZebaSpin Desalting columns with a molecular weight cut-off at 7 kDa (Thermo Fisher, #78606). Unlabeled DARPins, and sortase-A were removed by incubation with Ni2+-NTA magnetic beads for 15 minutes at room temperature. The absorption of the resulting DARPin solution was measured on a NanoPhotometer (Implen, NP80) at 280 nm. The concentration of DARPin was calculated as the ratio of absorption at 280 nm and the product of the DARPin-specific extinction coefficient and the length of the light path. The purity of labeled DARPins was verified using SDS-PAGE.

Bismaleimide-(polyethyleneglycol)3 (BM(PEG)3) was freshly diluted in dimethyl sulfoxide to reach 240 µM. Then, DARPin-GGGGGC and BM(PEG)3 were diluted in a 2.2:1 ratio in the phosphate buffer, reaching a final concentration of 26.4 µM for the DARPin and 12 µM BM(PEG)3. The solution was left under mild shaking for 4 hours at room temperature. Then, 5 µl fractions were stored at -80°C and thawed immediately before use.

### Reconstituted actin

G-actin stocks purchased from Hypermole were stored at -80°C after snap-freezing in liquid nitrogen. G-actin was diluted to reach 20 µM in 1xAPB (10xAPB: 20 mM Tris (pH 8.0), 500 mM KCl, 20 mM MgCl2, 10 mM Na2ATP) and 1xAB (10xAB: 250 mM imidazole in 1 M HCl (pH 7.4), 250 mM KCl, 40 mM MgCl2, 10 mM EGTA) buffer in H2O. The solution was left to polymerize for 15 minutes at room temperature. Optionally, phalloidin with a FITC or TRITC label was added with a final concentration of 0.2 mg/ml. F-actin was kept at 4°C for up to 2 weeks.

### F-actin cross-linking

A simple flow chamber was prepared by using double-sided sticky tape to separate two glass coverslips by 100-200 μm. Both coverslips were cleaned with isopropanol in an ultrasonic bath for a few seconds and dried under a fume hood. The bottom coverslip was additionally incubated with 1.0 ml of 0.1 mg/ml Poly-L-Lysine (PLL) (EMD Millipore Corp specialty media, A-005-C, LOT 3279047) for 30 minutes at room temperature, rinsed twice with ddH2O and dried overnight in a fume hood. After assembly, liquid was pipetted on the one opening and filled the chamber due to capillary forces. The chambers were rinsed with AB buffer just before adding F-actin.

ABdtt buffer was prepared freshly by mixing 100 µl of 10xAB buffer with 20 µl of 1M DTT in 880 µl H2O. Gas was removed under vacuum at 4°C for 15 minutes. F-actin was diluted to 3 µM in 50 µl ABdtt buffer and 1 µM TRITC-phalloidin was added. Additionally, either 1 µM fascin (Hypermole #8411-03) or 1µM covalent DARPin dimer 1ABD-1ABD was added and incubated for 1 h on ice. The mix was then diluted with minimal pipetting to 150 nM F-actin in ABdtt buffer and 15 µl were added to a flow chamber. F-actin was imaged on the plane directly above the PLL-coated glass surface using a laser scanning confocal microscope (Zeiss LSM880) with a Plan-Apochromat 63x/1.4 NA oil objective (Zeiss).

### Cell culture

The human osteosarcoma cell line U2OS was purchased from the DMSZ-German collection of microorganisms and cell cultures GmbH and was gifted from the laboratory of Kai Johnsson (Max Planck Institute for Medical Research, Heidelberg, Germany). U2OS cells were cultured as sub-confluent monolayers in McCoy 5A medium supplemented with 10% fetal bovine serum (FBS) and 100 U/ml penicillin-streptomycin at 37°C in a 5% CO2 incubator.

### Transient transfections

U2OS cells (8×10^4 cells) were seeded in 12-well culture plates for transient transfection. 24 hours after seeding, the cells reached 80% confluency and were transiently transfected using 800 ng plasmid DNA mixed with 1.6 µl lipofectamine 3000 and 1.6 µl P3000 reagent (Invitrogen) in 100 µl OptiMem according to the manufacturer’s protocol. To achieve co-transfections involving multiple plasmids, an equal weight of all plasmids was used to reach a total of 800 ng. Cells were harvested 24 hours after lipofection for further use.

### Optical Microscopy

Glass coverslips were coated with 10 µg/ml fibronectin (FN) (Sigma-Aldrich #F-1141) in phosphate-buffered saline (PBS) overnight at 4°C and subsequently washed three times with PBS. U2OS cells were seeded at a density of 4000 cells/cm²; after three hours, the medium was exchanged to CO2-independent medium (Gibco Biosciences #18045-054) with 10% FBS. Fluorescence images were acquired on a confocal laser scanning microscope (Zeiss LSM880) equipped with a Plan-Apochromat 63x/1.4 NA oil objective (Zeiss) and an incubation chamber set at 37°C.

### Optogenetic activation using an external LED light source

Light-triggered activation of the LOV2-SsrA domain on DARPins over the entire cell area was achieved by continuous illumination with a collimated 455 nm light-emitting diode (LED; Thorlabs #M455L4-C2) using a supply current of 0.35 A, unless noted otherwise. The LED was mounted at a 45 °C angle at a distance of approximately 15 cm from the sample. In this setup, the light intensity at the sample was 2.9 mW/cm² as measured with an integrating sphere photodiode power sensor (Thorlabs #S142C).

The association (activation) kinetics were quantified from time-lapse imaging during continuous LED illumination using NBD-mApple-SspB and 2ABD -SsrA.

The mean fluorescence intensity measured outside of a cell (background) was subtracted from the fluorescence intensity inside a region of interest (ROI) outlining part of a stress fiber and from the fluorescence intensity of the whole cell. Then, the ratio between the fluorescence intensity in the ROI and cell was calculated and normalized to the average fluorescence ratio before light illumination.

The data were fitted using a non-linear fit describing a one phase association 𝑦 = 𝑦*_0_* + (𝑦_𝑚_ −𝑦*_0_*)(*1* − 𝑒^−𝑘^(𝑡−𝑡0)) where *t_0_* = 9s and *y_0_* = 1.0 using GraphPad Prism Version 9.5.1.

For quantification of the dissociation dynamics the fluorescence intensity after iLID activation in a stress fiber was normalized to 1.0. The data were fitted using a non-linear fit model for a one phase decay 𝑦 = (𝑦*_0_* − 𝑦_𝑚_) ∗ 𝑒(−𝑘𝑡) + 𝑦_𝑚_ where 𝑦*_0_* = *1*.*0* using GraphPad Prism Version 9.5.1.

### Optogenetic activation of cells/subcellular regions using confocal microscopy

Activation of the iLID system in cells or subcellular ROIs was achieved by iterative scanning using the 458 nm Argon laser line of the LSM880 confocal microscope (Zeiss) during time-lapse image acquisition. For whole cell illumination, the 458 nm laser was activated during scanning. Illumination of a region corresponding to approximately half of a cell was achieved by scanning three times at a resolution of 0.1 µm/pixel with a speed of 0.58 µs/pixel in 120 s intervals. For subcellular activation, a smaller region of 81 µm² was scanned 20 times every 20 seconds at a resolution of 0.1 µm/pixel and a speed of 0.58 µs/pixel. The fluorescent DARPin-mApple-SspB constructs were concurrently monitored with a frame rate of 12 frames/minute. Image analysis was performed using custom Fiji (ImageJ 2.14.0) scripts to monitor the accumulation of DARPins inside the ROI. The mean fluorescence intensity in a region outside the cell was used for background subtraction. The resulting fluorescence intensities in the ROI and the whole cell were normalized to the fluorescence intensity in the respective region before light illumination.

In order to examine lamellipodia dynamics, a 9×9 μm² ROI at the cell edge was activated. Segmentation of the cell using the DARPin-mApple-SspB fluorescence signal was performed to calculate the difference in area between 0 and 5, 10 or 15 minutes within the ROI.

### Actin accumulation during optogenetic activation

Cells co-expressing Lifeact-HaloTag and the iLID-DARPin pair 1ABD-1ABD, or 2ABD-2ABD, were seeded on 10 µg/ml FN-coated glass-bottom dishes with cell culture medium containing 250 nM Halo-Ligand-Alexa647 conjugate. Three hours after cell seeding the medium was exchanged to CO2-independent medium and cells were transferred to a laser-scanning confocal microscope (Zeiss LSM880) with an environmental chamber set at 37°C. Fluorescence images of DARPin-mApple-SspB and Lifeact-Halo-Ligand-Alexa647 were acquired before and 20 minutes after subcellular activation of the iLID-system. Activation of the iLID-system in a ROI was performed as described above (Section Optogenetic activation of a subcellular region) for a duration of 20 minutes.

The background fluorescence intensity outside the cell was subtracted from the intensity in the ROI. The ratio of fluorescence intensity before and after iLID activation was calculated for DARPin-mApple-SspB and Lifeact-Halo-Ligand-Alexa647.

### Giant Unilamellar Vesicles (GUVs)

GUVs were produced by emulsion transfer. For this, the lipid-in-oil solution was prepared in 80% 1,2-dioleoyl-sn-glycero-3-phosphocholine (DOPC), 20% 1,2-dioleoyl-sn-glycero-3-phospho-(1′-rac-glycerol) (sodium salt) (DOPG) yielding a final concentration of 643 µM in 1 mL mineral oil. Optionally, the percentage of DOPC was reduced to add 0.5% Atto488 1,2-dioleoyl-sn-glycero-3-phosphoethanolamin (Atto488-DOPE) for visualization. The chloroform solvent of the lipids was removed in low-binding glass vials under vacuum for 20 minutes. The dried lipid film was re-hydrated in 60 µl n-decane, appended with 940 µl mineral oil and vortexed. The outside aqueous solution was prepared in water containing 5 mM imidazole-HCl at pH 7.6, and sucrose to adjust the osmolarity to 305 mOsm. 400 µl of the rehydrated lipids were added on top of 500 µl outside solution in a 2 ml eppendorf tube and left to form a lipid monolayer for 1 hour at room temperature. The inside aqueous solution was prepared in a volume of 10 µl, containing 20% Opti-Prep (5-[acetyl-[3-[acetyl-[3,5-bis(2,3-dihydroxypropylcarbamoyl)-2,4,6-triiodo-phenyl]amino]-2-hydroxy-propyl]amino]-N,N′-bis(2,3-dihydroxypropyl)-2,4,6-triiodo-benzene-1,3-dicarboxamide), 1x actin polymerization buffer, 5 µM G-actin and 2 µM covalent DARPin dimer 1ABD-1ABD in PBS. The inside solution was added to the remaining 100 µl of lipid-in-oil solution and pipetted up and down to create a water in oil emulsion. This was added on top of the oil and water layers and centrifuged at 380 rcf for 3 minutes. The upper oil film was carefully removed and the GUVs in the aqueous outside buffer were carefully pipetted with a cut pipette tip on glass coated with 5% BSA for observation under a confocal LSM880 microscope. Additionally, 3 µM SiR-actin was added to the aqueous outside buffer for F-actin visualization.

### Traction force microscopy

Polyacrylamide (PAA) hydrogels were fabricated based on the protocol from Christian *et al.*^42^ to serve as substrates for traction force microscopy (TFM). For hydrogels with an expected Young’s modulus of 16.7 kPa, a precursor solution was prepared by mixing 2.5 ml acrylamide (40% w/v), 0.75 ml bis-acrylamide (2% w/v), and 6.75 ml ddH2O^40^. The precursor solution was degassed for 30 minutes. The cross-linking reaction was initiated by addition of tetramethylethylenediamine (TEMED) and ammonium persulfate (APS), and a small volume of the reaction mixture was polymerized between two glass coverslips, one of which was functionalized with 3-(trimethoxysilyl)propyl methacrylate. Dark-red fluorescent beads, 200 nm in diameter (ThermoFisher Scientific, #F8807), were added to the precursor to serve as fiducial markers. After removing the top coverslip and washing the gel with PBS, the free surface of the hydrogel was coated with 10 µg/ml fibronectin in PBS through overnight incubation at 4°C in a humidified atmosphere. Finally, gels were washed twice with PBS before cell seeding.

Cells (transfected with iLiD pairs or non-transfected) were seeded on PAA hydrogels at a density of 3,000 cells/cm² in culture medium and left to adhere for 4 hours at 5% CO₂, 37 °C, before being washed once with PBS and once with culture medium. The cells on the hydrogels were then transferred to a Leica DMi8 epifluorescence microscope equipped with a 63×/1.4 objective and a 37 °C incubation chamber at 5% CO₂ for TFM imaging. Transfected cells were identified based on their fluorescence intensity stemming from the expression of the DARPin-mApple-SspB construct. First, baseline images under dark conditions were acquired. Transfected cells were then continuously illuminated using a 455 nm LED (Thorlabs #M455L4-C2) at 0.2 A for a duration of 15 minutes, after which another image of the beads was acquired. Finally, cells were detached from the substrate using 100 µl of 10% sodium dodecyl sulfate (SDS) and a reference image of the relaxed gel at the same spot was acquired. Datasets were processed using the particle image velocimetry (PIV) and Fourier transform traction cytometry (FTTC) plugins for ImageJ (v2.16.0/1.54p).^43^ Strain energy calculations were performed using Python (v3.9.6).

## Acknowledgements

We thank the Dieter Schwarz Foundation (DSS) for its generous support. The Max Planck Society and the Max Planck School Matter to Life are acknowledged for excellent support.

## Associated Content

Supporting Information

## Abbreviations

ABPs: actin-binding proteins
BM(PEG)3: Bismaleimide-(polyethyleneglycol)3
ABD: actin binding DARPin
BSA: bovine serum albumin
CI: confidence interval
DARPin: Designed Ankyrin Repeat Protein
DMEM: Dulbecco’s modified Eagle’s medium
DOPE: 1,2-dioleoylsn-glycero-3-phosphoethanolamin
DOPC: 1,2-dioleoyl-sn-glycero-3-phosphocholine
DOPG: 1,2-dioleoyl-sn-glycero-3-phospho-(1′-rac-glycerol)
F-actin: filamentous actin
FBS: fetal bovine serum
G-actin: globular actin
GUV: giant unilamellar vesicle
iLID: improved light-induced dimer
LED: light-emitting diode
LOV2: light-oxygen-voltage 2
mEGFP: monomeric enhanced green-fluorescent protein
NBD: non-binder DARPin
PAA: polyacrylamide
PBS: phosphate-buffered saline
PLL-PEG: poly-L-lysin-co-poly(ethylene glycol)
ROI: region of interest
SDS: sodium dodecyl sulfate
TFM: traction force microscopy
τ_1/2_: half-recovery time
UtrCH: utrophin calponin homology domain
U2OS: human bone osteosarcoma epithelial cells

## References

1. Lappalainen, P.; Kotila, T.; Jégou, A.; Romet-Lemonne, G., Biochemical and Mechanical Regulation of Actin Dynamics. Nat Rev Mol Cell Bio 2022, 23 (12), 836–852.

2. Pollard, T. D.; Goldman, R. D. J. C. S. H. p. i. b., Overview of the Cytoskeleton from an Evolutionary Perspective. 2018, 10 (7), a030288.

3. Clarke, D. N.; Martin, A. C. J. C. B., Actin-Based Force Generation and Cell Adhesion in Tissue Morphogenesis. 2021, 31 (10), R667–R680.

4. Jorgenson, T. D.; Baboolall, K. D.; Suarez, C.; Kovar, D. R.; Gardel, M. L.; Rowan, S. J., Highly Flexible Peg-Lifeact Constructs Act as Tunable Biomimetic Actin Crosslinkers. Soft Matter 2024, 20 (5), 971–977.

5. Lorenz, J. S.; Schnauß, J.; Glaser, M.; Sajfutdinow, M.; Schuldt, C.; Käs, J. A.; Smith, D. M., Synthetic Transient Crosslinks Program the Mechanics of Soft, Biopolymer-Based Materials. Adv Mater 2018, 30 (13), e1706092–e1706092.

6. Mulla, Y.; Avellaneda, M. J.; Roland, A.; Baldauf, L.; Jung, W.; Kim, T.; Tans, S. J.; Koenderink, G. H., Weak Catch Bonds Make Strong Networks. Nat Mater 2022, 21 (9), 1019–1023.

7. Effiong, U. M.; Khairandish, H.; Ramirez-Velez, I.; Wang, Y.; Belardi, B., Turn-on Protein Switches for Controlling Actin Binding in Cells. Nat Commun 2024, 15 (1), 5840.

8. Klewer, L.; Wu, Y. W., Light-Induced Dimerization Approaches to Control Cellular Processes. Chemistry–A European Journal 2019, 25 (54), 12452–12463.

9. Spiltoir, J. I.; Tucker, C. L., Photodimerization Systems for Regulating Protein-Protein Interactions with Light. Curr Opin Struc Biol 2019, 57, 1–8.

10. Kim, C.; Yun, S. R.; Lee, S. J.; Kim, S. O.; Lee, H.; Choi, J.; Kim, J. G.; Kim, T. W.; You, S.; Kosheleva, I.; Noh, T.; Baek, J.; Ihee, H., Structural Dynamics of Protein-Protein Association Involved in the Light-Induced Transition of Lov2 Protein. Nature Communications 2024, 15 (1).

11. Strickland, D.; Lin, Y.; Wagner, E.; Hope, C. M.; Zayner, J.; Antoniou, C.; Sosnick, T. R.; Weiss, E. L.; Glotzer, M., Tulips: Tunable, Light-Controlled Interacting Protein Tags for Cell Biology. Nature Methods 2012, 9 (4), 379–384.

12. Taslimi, A.; Zoltowski, B.; Miranda, J. G.; Pathak, G. P.; Hughes, R. M.; Tucker, C. L., Optimized Second-Generation Cry2–Cib Dimerizers and Photoactivatable Cre Recombinase. Nature Chemical Biology 2016, 12 (6), 425–430.

13. Guntas, G.; Hallett, R. A.; Zimmerman, S. P.; Williams, T.; Yumerefendi, H.; Bear, J. E.; Kuhlman, B., Engineering an Improved Light-Induced Dimer (Ilid) for Controlling the Localization and Activity of Signaling Proteins. Proceedings of the National Academy of Sciences 2015, 112 (1), 112–117.

14. Plückthun, A., Designed Ankyrin Repeat Proteins (Darpins): Binding Proteins for Research, Diagnostics, and Therapy. Annual Review of Pharmacology and Toxicology 2015, 55 (1), 489–511.

15. Bery, N.; Legg, S.; Debreczeni, J.; Breed, J.; Embrey, K.; Stubbs, C.; Kolasinska-Zwierz, P.; Barrett, N.; Marwood, R.; Watson, J. J. N. C., Kras-Specific Inhibition Using a Darpin Binding to a Site in the Allosteric Lobe. 2019, 10 (1), 2607.

16. Chabloz, A.; Schaefer, J. V.; Kozieradzki, I.; Cronin, S. J. F.; Strebinger, D.; Macaluso, F.; Wald, J.; Rabbitts, T. H.; Plückthun, A.; Marlovits, T. C.; Penninger, J. M., Salmonella-Based Platform for Efficient Delivery of Functional Binding Proteins to the Cytosol. Communications Biology 2020, 3 (1), 342.

17. Dreier, B.; Mikheeva, G.; Belousova, N.; Parizek, P.; Boczek, E.; Jelesarov, I.; Forrer, P.; Plückthun, A.; Krasnykh, V. J. J. o. m. b., Her2-Specific Multivalent ADARters Confer Designed Tropism to Adenovirus for Gene Targeting. 2011, 405 (2), 410–426.

18. Hartmann, J.; Münch, R. C.; Freiling, R.-T.; Schneider, I. C.; Dreier, B.; Samukange, W.; Koch, J.; Seeger, M. A.; Plückthun, A.; Buchholz, C. J. J. M. T.-M.; Development, C., A Library-Based Screening Strategy for the Identification of Darpins as Ligands for Receptor-Targeted Aav and Lentiviral Vectors. 2018, 10, 128–143.

19. Parizek, P.; Kummer, L.; Rube, P.; Prinz, A.; Herberg, F. W.; Plückthun, A. J. A. c. b., Designed Ankyrin Repeat Proteins (Darpins) as Novel Isoform-Specific Intracellular Inhibitors of C-Jun N-Terminal Kinases. 2012, 7 (8), 1356–1366.

20. Ivanova, J. R.; Benk, A. S.; Schaefer, J. V.; Dreier, B.; Hermann, L. O.; Plückthun, A.; Missirlis, D.; Spatz, J. P., Designed Ankyrin Repeat Proteins as Actin Labels of Distinct Cytoskeletal Structures in Living Cells. Acs Nano 2024, 18 (12), 8919–8933.

21. Fink, A.; Doll, C. R.; Relimpio, A. Y.; Dreher, Y.; Spatz, J. P.; Göpfrich, K.; Cavalcanti-Adam, E. A., Extracellular Cues Govern Shape and Cytoskeletal Organization in Giant Unilamellar Lipid Vesicles. Acs Synthetic Biology 2023, 12 (2), 369–374.

22. Weiss, M.; Frohnmayer, J. P.; Benk, L. T.; Haller, B.; Janiesch, J. W.; Heitkamp, T.; Börsch, M.; Lira, R. B.; Dimova, R.; Lipowsky, R.; Bodenschatz, E.; Baret, J. C.; Vidakovic-Koch, T.; Sundmacher, K.; Platzman, I.; Spatz, J. P., Sequential Bottom-up Assembly of Mechanically Stabilized Synthetic Cells by Microfluidics. Nat Mater 2018, 17 (1), 89-+.

23. Binz, H. K.; Stumpp, M. T.; Forrer, P.; Amstutz, P.; Plückthun, A. J. J. o. m. b., Designing Repeat Proteins: Well-Expressed, Soluble and Stable Proteins from Combinatorial Libraries of Consensus Ankyrin Repeat Proteins. 2003, 332 (2), 489–503.

24. Eitoku, T.; Nakasone, Y.; Matsuoka, D.; Tokutomi, S.; Terazima, M., Conformational Dynamics of Phototropin 2 Lov2 Domain with the Linker Upon Photoexcitation. J Am Chem Soc 2005, 127 (38), 13238–44.

25. Harper, S. M.; Neil, L. C.; Gardner, K. H., Structural Basis of a Phototropin Light Switch. Science 2003, 301 (5639), 1541–4.

26. Natwick, D. E.; Collins, S. R., Optimized Ilid Membrane Anchors for Local Optogenetic Protein Recruitment. ACS Synth Biol 2021, 10 (5), 1009–1023.

27. Hallett, R. A.; Zimmerman, S. P.; Yumerefendi, H.; Bear, J. E.; Kuhlman, B., Correlating in Vitro and in Vivo Activities of Light-Inducible Dimers: A Cellular Optogenetics Guide. ACS Synth Biol 2016, 5 (1), 53–64.

28. Park, J. S.; Burckhardt, C. J.; Lazcano, R.; Solis, L. M.; Isogai, T.; Li, L.; Chen, C. S.; Gao, B.; Minna, J. D.; Bachoo, R.; DeBerardinis, R. J.; Danuser, G., Mechanical Regulation of Glycolysis Via Cytoskeleton Architecture. Nature 2020, 578 (7796), 621–626.

29. Krause, M.; Gautreau, A., Steering Cell Migration: Lamellipodium Dynamics and the Regulation of Directional Persistence. Nat Rev Mol Cell Biol 2014, 15 (9), 577–90.

30. Mullins, R. D.; Heuser, J. A.; Pollard, T. D. J. P. o. t. N. A. o. S., The Interaction of Arp2/3 Complex with Actin: Nucleation, High Affinity Pointed End Capping, and Formation of Branching Networks of Filaments. 1998, 95 (11), 6181–6186.

31. Koestler, S. A.; Auinger, S.; Vinzenz, M.; Rottner, K.; Small, J. V. J. N. c. b., Differentially Oriented Populations of Actin Filaments Generated in Lamellipodia Collaborate in Pushing and Pausing at the Cell Front. 2008, 10 (3), 306–313.

32. Bieling, P.; Li, T. D.; Weichsel, J.; McGorty, R.; Jreij, P.; Huang, B.; Fletcher, D. A.; Mullins, R. D., Force Feedback Controls Motor Activity and Mechanical Properties of Self-Assembling Branched Actin Networks. Cell 2016, 164 (1-2), 115–127.

33. Doss, B. L.; Pan, M.; Gupta, M.; Grenci, G.; Mege, R. M.; Lim, C. T.; Sheetz, M. P.; Voituriez, R.; Ladoux, B., Cell Response to Substrate Rigidity Is Regulated by Active and Passive Cytoskeletal Stress. Proc Natl Acad Sci U S A 2020, 117 (23), 12817–12825.

34. Shao, H.; Wang, J. H.; Pollak, M. R.; Wells, A., Alpha-Actinin-4 Is Essential for Maintaining the Spreading, Motility and Contractility of Fibroblasts. Plos One 2010, 5 (11), e13921.

35. Ehrlicher, A. J.; Krishnan, R.; Guo, M.; Bidan, C. M.; Weitz, D. A.; Pollak, M. R., Alpha-Actinin Binding Kinetics Modulate Cellular Dynamics and Force Generation. Proc Natl Acad Sci U S A 2015, 112 (21), 6619–24.

36. Katsuta, H.; Okuda, S.; Nagayama, K.; Machiyama, H.; Kidoaki, S.; Kato, M.; Sokabe, M.; Miyata, T.; Hirata, H., Actin Crosslinking by Alpha-Actinin Averts Viscous Dissipation of Myosin Force Transmission in Stress Fibers. iScience 2023, 26 (3), 106090.

37. Katsuta, H.; Sokabe, M.; Hirata, H., From Stress Fiber to Focal Adhesion: A Role of Actin Crosslinkers in Force Transmission. Front Cell Dev Biol 2024, 12, 1444827.

38. Barai, A.; Mukherjee, A.; Das, A.; Saxena, N.; Sen, S., Alpha-Actinin-4 Drives Invasiveness by Regulating Myosin Iib Expression and Myosin Iia Localization. J Cell Sci 2021, 134 (23).

39. Wolfenson, H.; Meacci, G.; Liu, S.; Stachowiak, M. R.; Iskratsch, T.; Ghassemi, S.; Roca-Cusachs, P.; O’Shaughnessy, B.; Hone, J.; Sheetz, M. P., Tropomyosin Controls Sarcomere-Like Contractions for Rigidity Sensing and Suppressing Growth on Soft Matrices. Nat Cell Biol 2016, 18 (1), 33–42.

40. Tse, J. R.; Engler, A. J., Preparation of Hydrogel Substrates with Tunable Mechanical Properties. Curr Protoc Cell Biol 2010, *Chapter 10*, Unit 10 16.

41. Chen, I.; Dorr, B. M.; Liu, D. R., A General Strategy for the Evolution of Bond-Forming Enzymes Using Yeast Display. 2011, 108 (28), 11399–11404.

42. Christian, J.; Blumberg, J. W.; Probst, D.; Lo Giudice, C.; Sindt, S.; Selhuber-Unkel, C.; Schwarz, U. S.; Cavalcanti-Adam, E. A., Control of Cell Adhesion Using Hydrogel Patterning Techniques for Applications in Traction Force Microscopy. J Vis Exp 2022, (179).

43. Tseng, Q.; Duchemin-Pelletier, E.; Deshiere, A.; Balland, M.; Guillou, H.; Filhol, O.; Thery, M., Spatial Organization of the Extracellular Matrix Regulates Cell-Cell Junction Positioning. Proc Natl Acad Sci U S A 2012, 109 (5), 1506–11.

