## Supplementary Figures S1-S3 for "Optogenetic cross-linking of the actin cytoskeleton using DARPins"

### Optogenetic regulation of the cytoskeleton using DARPin-based actin cross-linkers

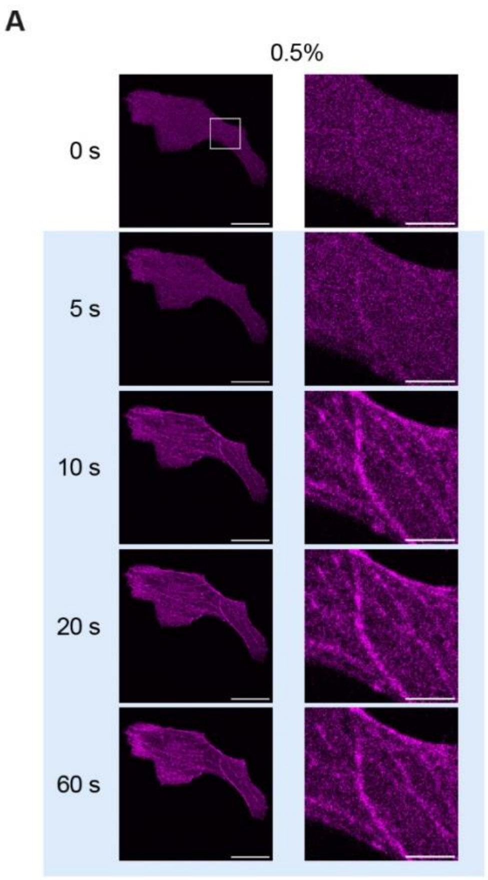


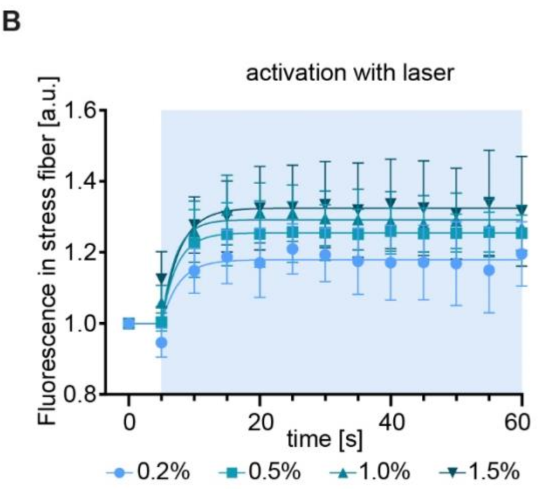


C

| laser power | association $\tau_{1/2}$ | 95% CI |
|---|---|---|
| 0.2% | 1.8 s | <11.8 s |
| 0.5% | 1.6 s | <5.8 s |
| 1.0% | 1.5 s | <4.7 s |
| 1.5% | 1.9 s | <8.9 s |

**Figure S1. Activation dynamics with a 458 nm laser light. (A)** Live U2OS cells co-expressing 2ABD-SsrA and NBD-mApple-SspB (magenta) were imaged with 5 s intervals by confocal scanning. Immediately after t= 0 s the 458 nm laser light of 0.5% power was switched on for the following image acquisitions illuminating the entire cells. Details of the area marked with a white rectangle are shown next to the whole cell images (scale bars whole cell: 20 µm, scale bars zoom: 5 µm). **(B)** Fluorescence intensity increase in a stress fiber upon illumination of the whole cell with 458 nm laser light at different laser powers in 5 s intervals. Illumination began after 5 s (blue rectangle in background). **(C)** The association half-times for different laser powers are listed.

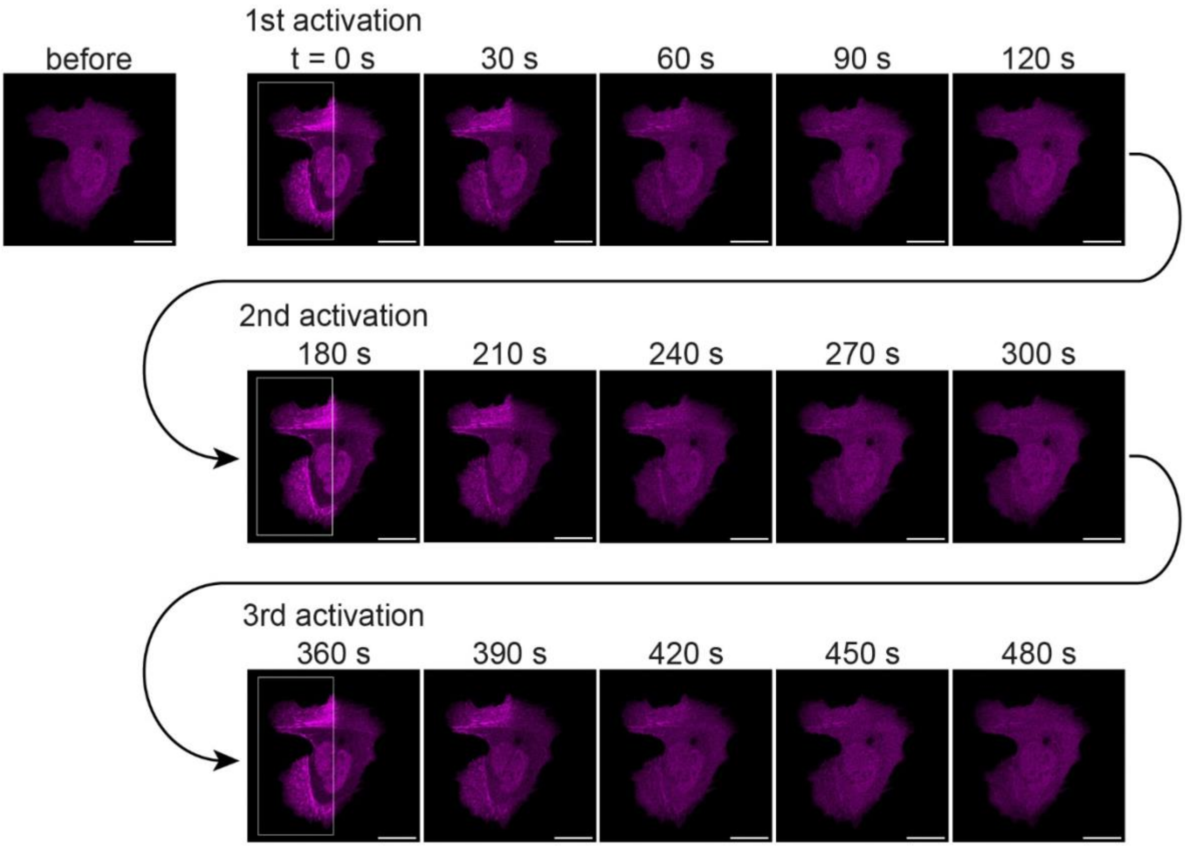


**Figure S2. Repeated, reversible and subcellular activation with 458 nm laser light.** Live U2OS cells co-expressing 2ABD-SsrA and NBD-mApple-SspB (magenta). The area in the white rectangle was scanned with 458 nm laser light in 120 s intervals for three repetitions. At 0 s, 180 s and 360 s the cell was imaged immediately after iLID activation in the rectangle (scale bars = 20 µm).

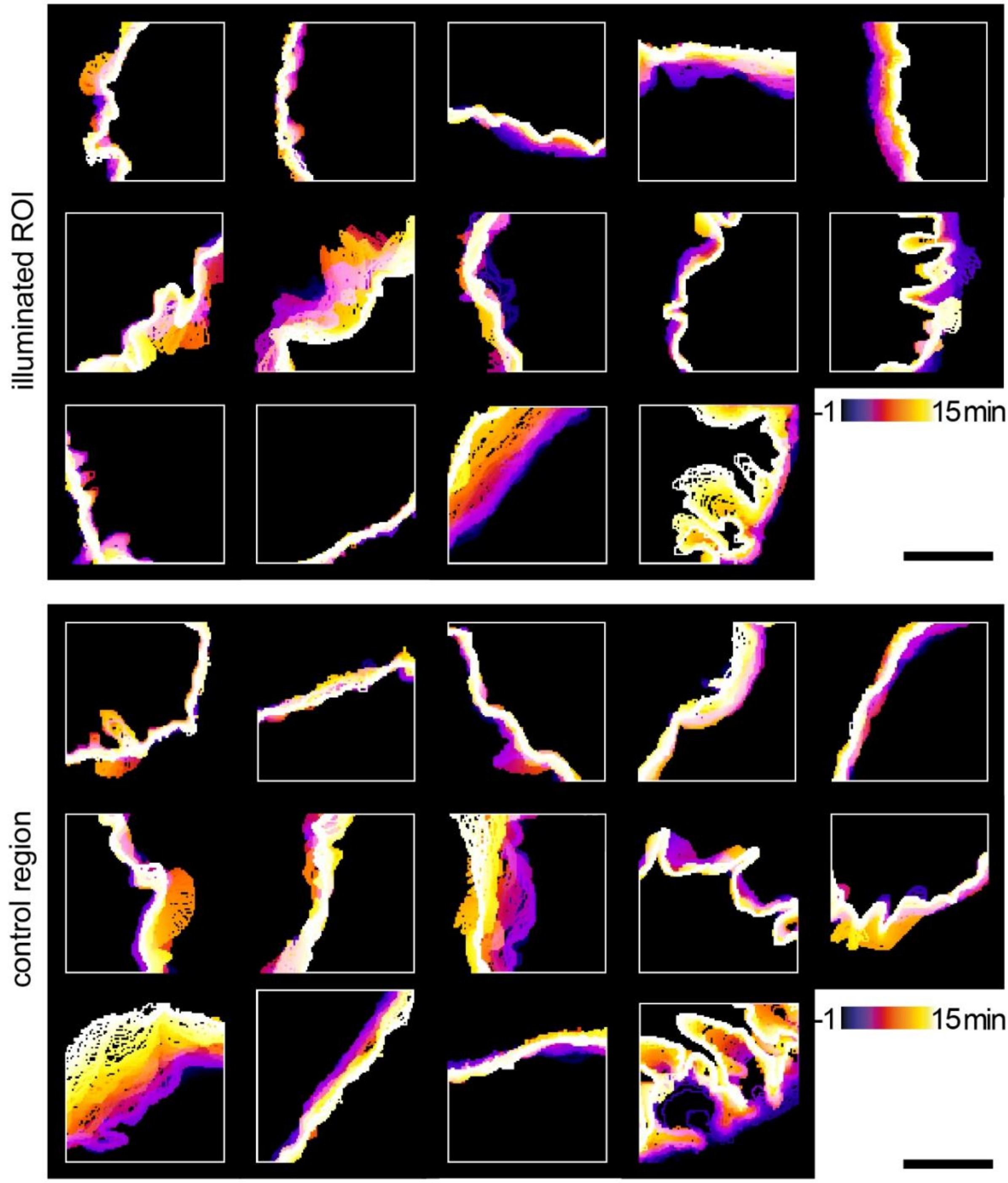


**Figure S3. Cell outlines in regions including lamellipodia.** Actin was crosslinked in ROIs including the lamellipodia region of U2OS cells transfected with iLID-DARPin pair 1ABD-1ABD for 15 min after one initial minute without crosslinking. The outline of the cell within the ROI was determined every 5 s and the outlines were plotted with a temporal color code (from blue to white). Non-crosslinked control regions in the same cells that also include the lamellipodium were analysed the same way (scale bars = 5 µm; n=14, N=3).
